# DHCR7 as a novel regulator of ferroptosis in hepatocytes

**DOI:** 10.1101/2022.06.15.496212

**Authors:** Naoya Yamada, Tadayoshi Karasawa, Takanori Komada, Takayoshi Matsumura, Chintogtokh Baatarjav, Junya Ito, Kiyotaka Nakagawa, Daisuke Yamamuro, Shun Ishibashi, Kouichi Miura, Naohiro Sata, Masafumi Takahashi

## Abstract

Recent evidence indicates that ferroptosis is implicated in the pathophysiology of various liver diseases; however, the mechanism of ferroptosis regulation in the liver is poorly understood. Here, using the whole-genome screening approach, we identified 7-dehydrocholesterol reductase (DHCR7), the terminal enzyme of cholesterol biosynthesis, as a novel regulator of ferroptosis in hepatocytes. Genetic and pharmacological inhibition (with AY9944) of DHCR7 suppressed lipid peroxidation and ferroptosis in human hepatocellular carcinoma Huh-7 cells. DHCR7 inhibition increased its substrate, 7-dehydrocholesterol (7-DHC), and extrinsic 7-DHC supplementation in turn suppressed ferroptosis. On the other hand, cholesterol deprivation had no effect on ferroptosis. A 7-DHC-derived oxysterol metabolite, 3β,5α-dihydroxycholest-7-en-6-one (DHCEO), was increased by a ferroptosis inducer RSL-3 in DHCR7-deficient cells, suggesting that the ferroptosis-suppressive effect of DHCR7 inhibition was driven by intracellular 7-DHC as a radical scavenger. While extrinsic 7-DHC supplementation suppressed ferroptosis in various cancer cells, pharmacological DHCR7 inhibition by AY9944 showed cell-type specific effects, which could be explained by high *DHCR7* expression in Huh-7 cells. We further showed that AY9944 suppressed ferroptosis in murine primary hepatocytes *in vitro* and systemic administration of AY9944 inhibited hepatic ischemia-reperfusion injury *in vivo*. These findings provide new insights into the regulatory mechanism of liver ferroptosis and suggest that DHCR7 inhibition is a potential therapeutic option for ferroptosis-related liver diseases.

## Introduction

Iron-dependent cell death, ferroptosis, was originally identified as a new form of regulated cell death in RAS-mutated cancer cells. It occurs when intracellular glutathione peroxidase 4 (GPX4) is inhibited directly or indirectly by a decrease in glutathione (GSH) levels (1, 2). GPX4 inhibition accumulates iron-dependent lipid peroxides derived from polyunsaturated fatty acid (PUFA)-containing phospholipids, leading to cellular/subcellular membrane damage and eventually cell death without specific effector molecules. Although ferroptosis was initially discovered in cancer cells, recent research has revealed that ferroptosis is implicated in the pathogenesis of various human diseases. We and other investigators have demonstrated that ferroptosis plays a crucial role in the development of acute and chronic liver diseases, including acetaminophen-induced acute liver failure, hepatic ischemia-reperfusion injury (IRI), hemochromatosis, non-alcoholic steatohepatitis (NASH), and alcoholic liver disease (ALD) (3–6). However, the regulatory mechanism of ferroptosis in the liver remains less understood. Furthermore, since the liver is a central organ for GSH synthesis and lipid and iron metabolism (6), which are the key factors to engaging ferroptosis, we postulated an unknown mechanism of ferroptosis regulation in the liver. Accordingly, we performed CRISPR/Cas9-mediated whole-genome screening and identified 7-dehydrocholesterol reductase (DHCR7), the terminal enzyme of cholesterol biosynthesis, as a novel regulator of ferroptosis in hepatocytes. Ferroptosis in human hepatocellular carcinoma-derived cells was suppressed by genetic and pharmacological inhibition of DHCR7 and also by extrinsic supplementation with the substrate for DHCR7, 7-dehydrocholesterol (7-DHC). Furthermore, DHCR7 inhibition suppressed ferroptosis in murine primary hepatocytes *in vitro* and inhibited hepatic IRI *in vivo*. Our findings reveal a novel regulatory mechanism of hepatic ferroptosis and highlight DHCR7 and 7-DHC as potential therapeutic targets for ferroptosis-related liver diseases.

## Materials and Methods

### Cell culture and reagents

Huh-7 (human hepatocellular carcinoma), OVISE (human ovarian clear cell carcinoma), and HT-1080 (human fibrosarcoma) cells were obtained from the Japanese Collection of Research Biosources (JCRB) Cell Bank (Japan). PLC/PRF/5 and HLE (human hepatocellular carcinoma) cells were obtained from The Health Science Research Resource Bank (HSRRB, Japan). SKHep1 (human hepatocellular carcinoma) and Panc-1 (human pancreatic ducal adenocarcinoma) cells were obtained from ATCC. The cells were cultured in Dulbecco’s modified Eagle’s medium (DMEM; Wako, Osaka, Japan) supplemented with 10% fetal calf serum (FCS; Dainippon Pharmaceutical Company, Osaka, Japan) and antibiotics. LentiX293T (Takara Bio, Shiga, Japan) cells were cultured in DMEM supplemented with 10% FCS and 1 mM sodium pyruvate. Primary cultured hepatocytes were isolated from 8-week-old male C57BL/6 mice as previously reported (7). Isolated hepatocytes were seeded at 1×10^4^ cells/well on 96-well plates and cultured in DMEM with 10% FCS for 2 h, followed by serum-free DMEM medium overnight.

Ferrostatin-1 (Fer-1, #17729) and AY9944 were purchased from Cayman Chemical (Ann Arbor, MI). Deferoxamine (DFO, #120727), RSL-3, and 7-dehydrocholesterol (7-DHC) were purchased from Abcam (Cambridge, MA), AdooQ Bioscience (Irvine, CA), and AdipoGen (Farmingdale, NY), respectively. Methyl-β-cyclodextrin (Mβ-CD) and other reagents were obtained from Sigma-Aldrich (St. Louis, MO) unless otherwise specified. Fer-1, RSL-3, and 7-DHC were dissolved in dimethyl sulfoxide (DMSO), and other reagents were dissolved in PBS. In *in vitro* experiments, cells were treated with Fer-1, AY9944, 7-DHC, and Mβ-CD 1 h prior to RSL-3 treatment, unless otherwise specified.

### Lentiviral preparation

LentiX293T cells were co-transfected together with LentiCRISPRv2, pLP1, pLP2, and pVSVG using PEI MAX (Polysciences, Warrington, PA, USA) to prepare the lentiviral vectors. Culture media containing the lentiviral vectors were collected 3 days after transfection. The collected media were filtered with a 0.45-μm filter and ultracentrifuged at 21,000 rpm using a SW55 Ti rotor (Beckman Coulter, Brea, CA, USA), and the pellets were resuspended in PBS containing 5% FCS. The lentivirus titer was measured using a Lentivirus qPCR Titer kit (Applied Biological Materials, Richmond, BC, Canada).

### Genome-wide CRISPR screening

Genome-wide CRISPR screening was performed using the Human GeCKOv2 CRISPR knockout pooled library (Addgene #1000000048). For sgRNA library transduction, 1 × 10^7^ Huh7 cells were infected with GeCKOv2 library virus at a target MOI of 0.3 in the presence of 8 μg/mL polybrene. Infected cells were selected by 2 μg/mL of puromycin for 3 days. The positively selected cells were expanded and treated with 0.1 μM RSL3 in the presence or absence of 50 mM linoleic acid (LA) and 0.03 μM RSL3 for 24 h. The surviving cells were further expanded and selected by repeated stimulation for 3 times and 5 times, separately. After selection, the genomic DNA was extracted from the surviving cells with phenol-chloroform. The sgRNA cassette was PCR-amplified and barcoded with sequencing adaptor using KOD One PCR Master Mix (TOYOBO). The PCR products were purified with a GEL/PCR purification column (FAVORGEN), quantified by Quant-iT PicoGreen dsDNA reagents and kits (Thermo Fisher), and sequenced on a Miseq sequencer (Illumina), loaded at 30 % spike-in of PhiX DNA. The sequence data were processed with the MAGeCK algorithm (8).

### CRISPR/Cas9-mediated genome editing

*DHCR7* and *GPX4* genes were mutated by CRISPR/Cas9 in Huh-7 cells. The sgRNA targeting each gene was designed with CRISPR direct (http://crispr.dbcls.jp) and is listed in Supplementary Table 1. The sgRNAs were subcloned into LentiCRISPRv2, which was a gift from Feng Zhang (Addgene plasmid #52961; http://n2t.net/addgene: 52961; RRID: Addgene_52961). For lentiviral transduction, Huh-7 cells were incubated with lentiviral vectors for 16 h in the presence of 8 μg/mL polybrene. The transduced cells were selected by incubating them with 2 μg/mL puromycin for at least 2 days. *GPX4*-KO cells were maintained in DMEM with 10% FCS in the presence of a low dose (0.1 μM) of Fer-1.

### Cell death assay

Cytotoxicity was determined as lactate dehydrogenase (LDH) activity in cultured supernatants using a cytotoxicity detection kit (#11644793001; Roche, Mannheim, Germany), according to the manufacturer’s instructions. To determine the total cellular LDH activity, cells were lysed with 2% TritonX100, and L-lactate dehydrogenase from rabbit muscle (Roche) was used as a standard. Cell death was also assessed by SYTOX Green (#S7020; Thermo Fisher Scientific), a membrane-impermeable DNA dye that enters dead cells. Nuclei were co-stained with Hoechst 33342 (No. 346-07951, Dojindo, Kumamoto, Japan). Images were captured by using confocal microscopy (FLUOVIEW FV10i; Olympus, Tokyo, Japan). Cell viability was determined by the MTT [3-(4,5-di-methylthiazol-2-yl)-2,5-diphenyltetrazolium bromide] (Invitrogen, Waltham, MA) reduction assay. Cytotoxicity or cell viability was evaluated 24 h after RSL-3 treatment or 48 h after Fer-1 withdrawal in *GPX4-knockout* (KO) Huh7 cells.

### Assessment of lipid peroxidation

Cells were cultured overnight and labeled with 5 μM C11-BODIPY^581/591^ (#D3861; Thermo Fisher Scientific) in 0.1% BSA/DMEM for 1 h before RSL-3 treatment. Fluorescence intensity was measured by using a multimode microplate reader (Spark, TECAN). Nuclei were labeled with Hoechst 33342 (#346-07951, Dojindo, Kumamoto, Japan). Representative images were obtained at 3 h after RSL-3 treatment using an FV10i confocal laser scanning microscope (Olympus, Tokyo).

### Real-time reverse transcription-polymerase chain reaction (RT-PCR)

Total RNA was prepared using ISOGEN (Nippon Gene Co., Ltd., Toyama, Japan) according to the manufacturer’s instructions. Real-time RT-PCR analysis was performed using the Thermal Cycler Dice Real-Time System II (Takara Bio Inc., Shiga, Japan) to detect the mRNA expression of *GPX4, SLC7A11, ACSL4, AIFM2, DHODH, DHCR7*, and *ACTB*. The primers are listed in Supplementary Table 2. The expression levels of each target gene were normalized by subtracting the corresponding β-actin threshold cycle (CT) value; normalization was carried out using the ΔΔCT comparative method.

### LC-MS/MS analysis for 7-DHC and DHCEO

Intracellular 7-DHC and DHCEO levels were analyzed by liquid chromatograph-mass spectrometry (LC-MS/MS). The detailed protocol is described in the supplementary file.

### Animal experiments

All experiments in this study were performed in accordance with the Jichi Medical University Guide for Laboratory Animals (Permit Nos. 17141-02 and 20107-02). C57BL/6J mice were purchased from SLC Japan (Shizuoka, Japan). Mice were housed (4/cage, RAIR HD ventilated Micro-Isolator Animal Housing Systems, Lab Products, Seaford, DE) in an environment maintained at 23 ± 2°C with *ad libitum* access to food and water under a 12-h light/dark cycle with lights on from 8:00 to 20:00. Partial hepatic ischemia was produced as previously described with some modifications (9, 10). Briefly, mice were anesthetized with isoflurane. Midline laparotomy was performed and an atraumatic clip (Fine Science Tools, Foster City, CA) was placed across the portal vein, hepatic artery, and bile duct to interrupt blood supply to the left lateral and median lobes (~70%) of the liver. After 60 min of partial hepatic ischemia, the clip was removed to initiate reperfusion. Sham control mice underwent the same protocol without vascular occlusion. Mice were sacrificed 3 h after reperfusion, and samples of blood and ischemic lobes were collected. AY9944 (25 mg/kg) or vehicle (PBS) was administered intraperitonially 1 h prior to open laparotomy. Serum levels of AST and ALT were measured using a Fuji-DRYCHEM chemical analyzer (Fuji Film, Tokyo, Japan) according to the manufacturer’s instructions. The paraffin-embedded tissue sections were stained with hematoxylin and eosin (HE). The severity of liver injury was graded according to a previous report (11).

### Statistics

Data are expressed as the mean ± standard error of the mean (SEM). Differences between two groups were determined by Mann-Whitney’s U test. Differences between multiple groups were determined by one-way analysis of variance (ANOVA) or two-way ANOVA combined with Tukey’s post hoc test. All analyses were performed using GraphPad Prism version 7 (San Diego, CA). A *p* value of < 0.05 was considered to be statistically significant.

## Results

### Genetic deletion of DHCR7 suppresses lipid peroxidation and ferroptosis in human hepatocellular carcinoma Huh-7 cells

To identify unknown regulators of ferroptosis in the liver, we performed CRISPR/Cas9-mediated whole-genome screening in human hepatocellular carcinoma Huh-7 cells. The cells were transduced with a lentiviral sgRNA library and treated with a lethal dose of the ferroptosis inducer RSL-3 in the presence or absence of LA, which further promotes ferroptosis (12) (Fig. 1A). After selecting ferroptosis-resistant cells four times, we analyzed enriched gRNAs in surviving cells and revealed that gRNAs targeting *DHCR7* were markedly enriched in both RSL-3- and RSL-3/LA-resistant cells (34.2% of all sequenced gRNAs) (Fig. 1B and C). To validate the screening, we generated *DHCR7*-KO-Huh-7 cells (Fig. S1) and confirmed that *DHCR7*-KO cells were resistant to RSL-3-induced ferroptosis (Fig. 1D and E). Lipid peroxidation, a hallmark of ferroptosis, was also suppressed in *DHCR7*-KO cells (Fig. 1F, Fig. S2A). The mRNA expression of major ferroptosis-regulating molecules such as *GPX4, SLC7A11, ACSL4, AIFM2*, and *DHODH*, was not influenced by *DHCR7* deficiency (Fig. 1G), suggesting that DHCR7 is a novel ferroptosis regulator driven by a mechanism that is distinct from previously described pathways.

**Figure 1.**
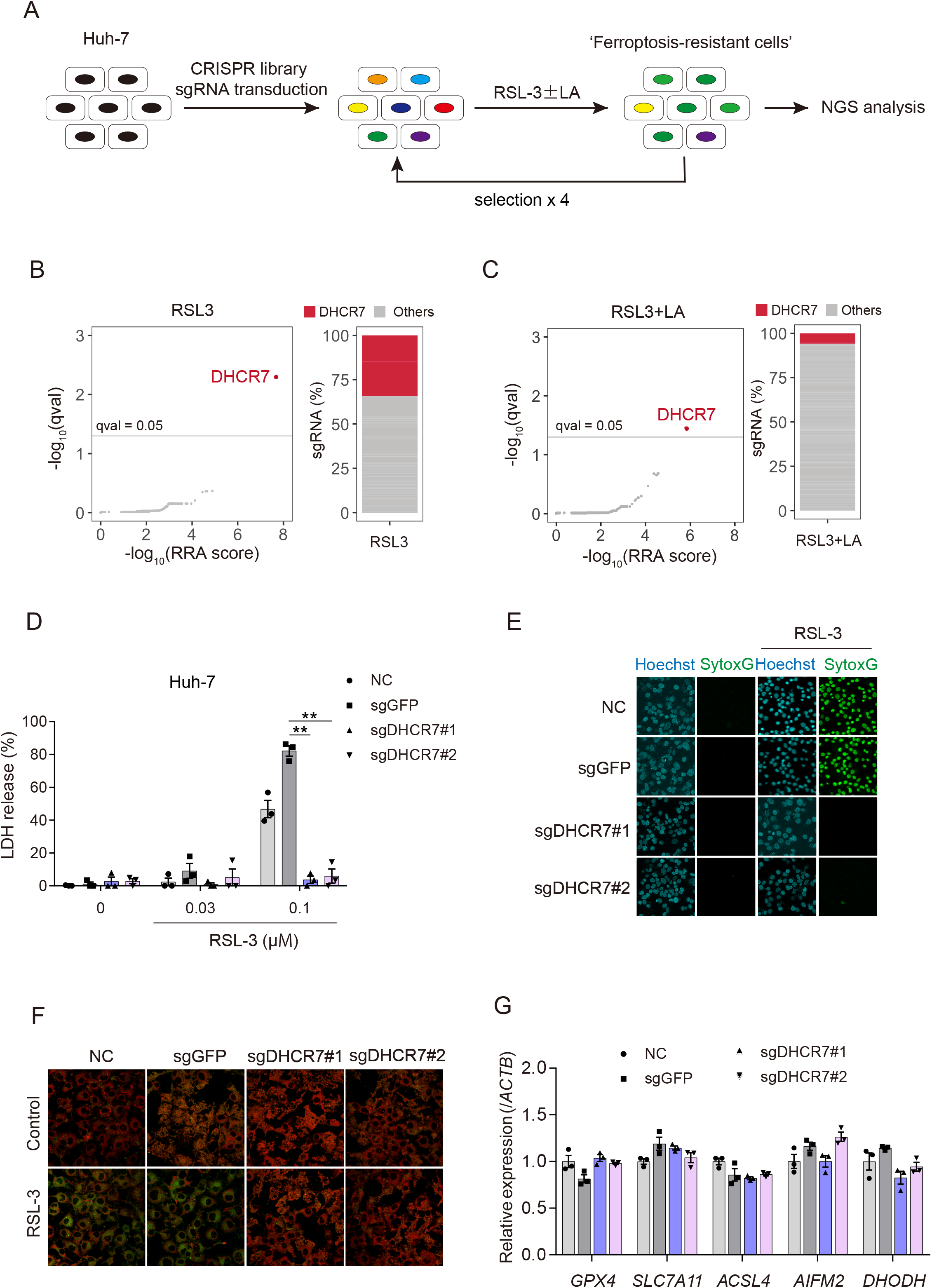
Genetic deletion of DHCR7 suppresses lipid peroxidation and ferroptosis in Huh-7 cells. (A) Strategy of CRISPR/Cas9-mediated whole-genome screening. Huh-7 cells were treated with RSL-3 (0.1 μM) or RSL-3 (0.03 μM) with LA (50 mM) for 24 h. The ferroptosis-resistant cells were selected 4 times, and the genomic DNA was purified and analyzed by NGS. (B and C) DHCR7 gRNAs were enriched in ferroptosis-resistant cells. (D and E) Control (sgGFP) and *DHCR7*-KO (sgDHCR7) Huh-7 cells were treated with or without RSL-3 (0.03 and 0.1 μM) for 24 h. Cytotoxicity and cell death were assessed by an LDH release assay and SYTOX Green, respectively. (F) Lipid peroxidation was assessed by C11-BODIPY^581/591^ staining. (G) The mRNA levels of *GPX4, SLC7A11, ACSL4, AIFM2*, and *DHODH* were assessed by real-time RT-PCR analysis. Statistical significance was calculated using two-way ANOVA (D) or one-way ANOVA (G) with Tukey’s post hoc test. Data are expressed as dot plots and means ± SEM. \**p* < 0.05, \*\**p* < 0.01.

### Pharmacological inhibition of DHCR7 ameliorates ferroptosis

DHCR7 is an enzyme that acts in the final step of the cholesterol biosynthesis pathway and converts 7-DHC to cholesterol (Fig. 2A). We used the DHCR7 inhibitor AY9944 and confirmed that AY9944 suppressed RSL-3-induced ferroptosis in Huh-7 cells in a dose-dependent manner (Fig. 2B and C, Fig. S2B). To exclude the possibility that the effect was limited to RSL-3-induced ferroptosis, we examined another type of ferroptosis. *GPX4*-KO cells were maintained with a minimum dose of Fer-1, and underwent ferroptosis at 48 h after Fer-1 withdrawal (termed “spontaneous ferroptosis”). Fer-1, the iron chelator DFO, and AY9944 suppressed this spontaneous ferroptosis (Fig. 2D), indicating that the suppressive effect of AY9944 is not limited to RSL-3-induced ferroptosis.

**Figure 2.**
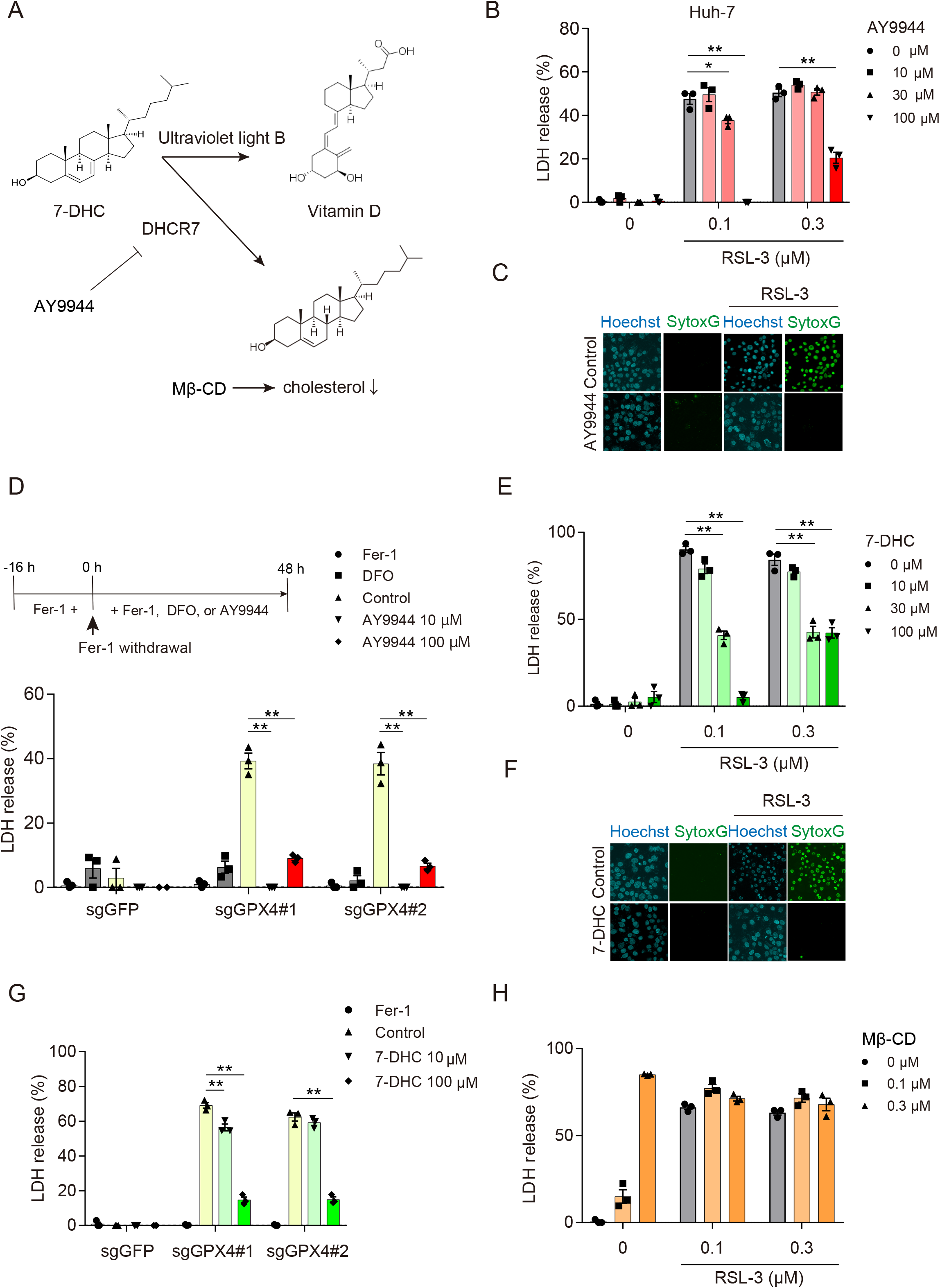
Pharmacological DHCR7 inhibition and 7-DHC supplementation ameliorates ferroptosis. (A) DHCR7 metabolic pathway in cholesterol synthesis. (B and C) Huh-7 cells were pretreated with AY9944 (0–100 μM) for 1 h, and then treated with RSL-3 for 24h. Cytotoxicity and cell death were assessed by an LDH release assay and SYTOX Green, respectively. (D) Control (sgGFP) and *GPX4*-KO (sgGPX4) cells were pretreated with Fer-1 (0.5 μM), DFO (100 μM), or AY9944 (10 and 100 μM) for 1 h, followed by Fer-1 withdrawal. Cytotoxicity at 48 h was assessed by an LDH release assay. (E and F) Cells were pretreated with 7-DHC for 1 h, and then treated with RSL-3 for 24 h. Cytotoxicity and cell death were assessed by an LDH release assay and SYTOX Green, respectively. (G) *GPX4*-KO cells were pretreated with 7-DHC for 1 h, followed by Fer-1 withdrawal. Cytotoxicity at 48 h was assessed by an LDH release assay. (H) Cells were pretreated with Mβ-CD (0.1 μM) for 1 h, and then treated with RSL-3 for 48 h. Cytotoxicity was assessed by LDH release assay. Statistical significance was calculated using the two-way ANOVA with Tukey’s post hoc test. Data are expressed as dot plots and means ± SEM. \**p* < 0.05, \*\**p* < 0.01.

### 7-DHC accumulation, but not cholesterol deprivation, prevents ferroptosis

Since DHCR7 converts 7-DHC to cholesterol (13), DHCR7 inhibition leads to intracellular accumulation of 7-DHC and loss of intracellular cholesterol (14). To elucidate which primarily contributed to ferroptosis suppression, we compared the effect of extrinsic 7-DHC supplementation and cholesterol deprivation. Interestingly, extrinsic 7-DHC supplementation suppressed both RSL-3-induced and spontaneous ferroptosis in a dose-dependent manner (Fig. 2E–G). On the contrary, cholesterol depletion by the cholesterol-extracting agent Mβ-CD failed to suppress ferroptosis (Fig. 2H). Taken together, these findings suggest that the accumulation of 7-DHC, but not cholesterol deprivation, is the main mechanism by which DHCR7 inhibition suppresses ferroptosis.

### 7-DHC is increased by DHCR7 inhibition and converted to oxysterol metabolites

To further investigate the ferroptosis-suppressive mechanism of DHCR7 inhibition through 7-DHC accumulation, we performed LC-MS/MS analysis and found that intracellular 7-DHC levels in *DHCR7*-KO cells were much higher than those in control cells (Fig. 3A). AY9944-treated cells also showed a moderate increase of intracellular 7-DHC (Fig. 3B). Notably, the elevation of 7-DHC levels in AY9944-treated cells was diminished by RSL-3; therefore, we assumed that 7-DHC could be metabolized to other forms by RSL-3. 7-DHC is known for its highly oxidizable property and its modification to various kinds of oxysterol metabolites (15). In particular, 7-DHC has been reported to easily react with peroxy radicals, compared to arachidonic acid, oxidation of which is critical for ferroptosis (16). Hence, we hypothesized that 7-DHC is oxidized and functions as a radical scavenger to prevent ferroptosis (15, 17). To test this hypothesis, we performed LC-MS/MS analysis to assess DHCEO, a major 7-DHC-derived oxysterol metabolite (15, 17), in RSL-3-treated and untreated *DHCR7*-KO cells. As expected, DHCEO was detected only in *DHCR7*-KO cells, and was significantly increased by RSL-3 (Fig. 3C and D), suggesting that 7-DHC is primarily oxidized instead of membrane phospholipids and restrains ferroptosis.

**Figure 3.**
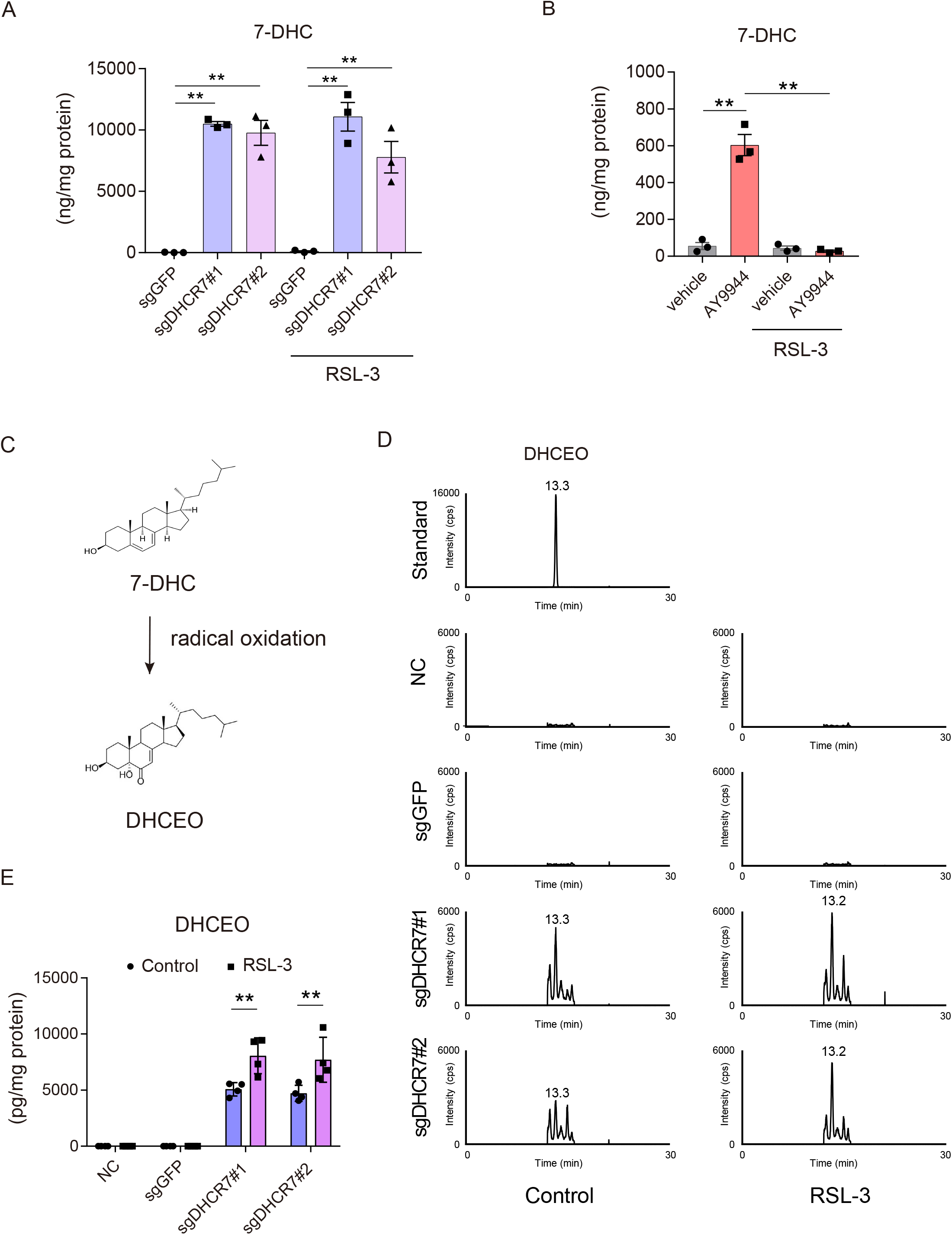
7-DHC is increased by DHCR7 inhibition and converted to oxysterol metabolites. (A) Control (sgGFP) and *DHCR7*-KO (sgDHCR7) Huh-7 cells were treated with or without RSL-3 (0.1 μM) for 16 h. (B) Huh-7 cells were pretreated with AY9944 (100 μM) for 1 h, and then treated with RSL-3 for 16 h. (A and B) Intracellular 7-DHC levels were assessed using LC-MS/MS analysis. (C) DHCEO is a major radical oxidation product of 7-DHC. (D and E) Control (sgGFP) and *DHCR7*-KO (sgDHCR7) Huh-7 cells were treated with or without RSL-3 (0.1 μM) for 16 h. Intracellular DHCEO levels were assessed using LC/MS-MS analysis. Statistical significance was calculated using the two-way ANOVA with Tukey’s post hoc test.

### While 7-DHC is a robust and cell-type-independent ferroptosis inhibitor, the effect of AY9944 is dependent on DHCR7 expression

Next, we studied the effects of 7-DHC and AY9944 on cells from various organs. 7-DHC almost completely suppressed ferroptosis in ferroptosis-sensitive cancer cells: human fibrosarcoma HT-1080, human pancreatic ducal adenocarcinoma Panc-1, and human ovarian clear cell carcinoma OVISE cells (Fig. 4A). On the other hand, unexpectedly, AY9944 had little effects on ferroptosis in these cells (Fig. 4A and B). These findings prompted us to analyze the expression levels of *DHCR7* in various cancer cells. A gene expression database of cancer cell lines (CellExpress, http://cellexpress.cgm.ntu.edu.tw) revealed that among various kinds of cancer cell lines, *DHCR7* expression in liver cancer cells was higher than those in most cancer cells of other origins (Fig. 4C) (18). Moreover, the *DHCR7* expression in Huh-7 cells was not only the highest among liver cancer cells but also remarkably higher than those in other ferroptosis-sensitive cancer cells including HT-1080, Panc-1, and OVISE cells (Fig. 4D). Therefore, we speculated that AY9944 could be effective only in cells with high *DHCR7* expression. To prove this, we examined several human hepatocellular carcinoma-derived cells with low *DHCR7* expression. As expected, AY9944 showed no effects on RSL-3-induced ferroptosis in these cells (Fig. 4E). Collectively, these results revealed that 7-DHC supplementation inhibits ferroptosis regardless of cell type, whereas AY9944 is a cell-type-specific ferroptosis inhibitor, the activity of which is likely dependent on the enzymatic activity of DHCR7.

**Figure 4.**
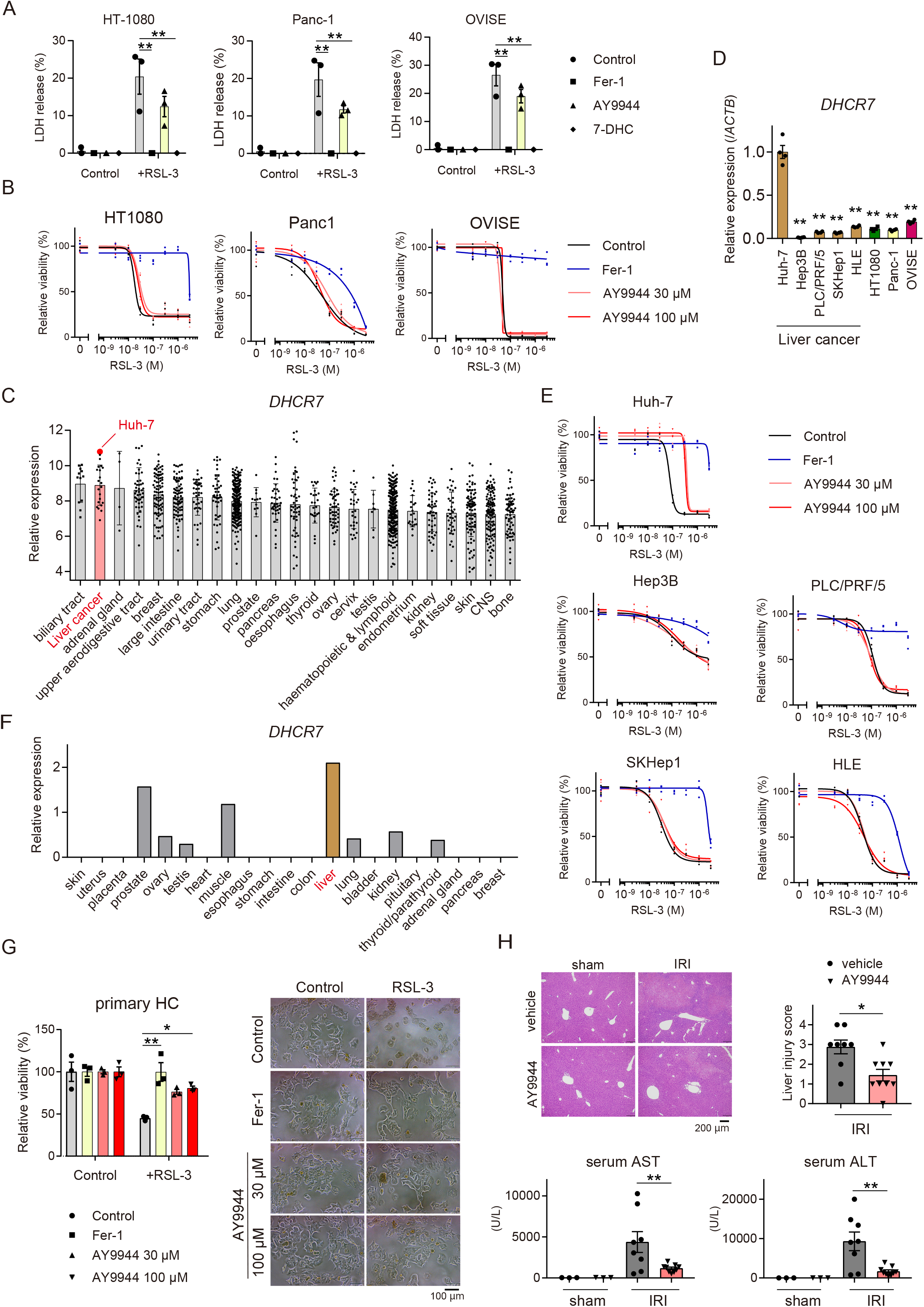
DHCR7 is a hepatocyte-specific regulator of ferroptosis. (A) HT-1080, Panc-1, and OVISE cells were pretreated with Fer-1 (1 μM), AY9944 (100 μM), or 7-DHC (100 nM), and then treated with RSL-3 (1 μM) for 24 h. Cytotoxicity was analyzed by an LDH release assay. (B) Cells were pretreated with Fer-1 (0.5 μM) or AY9944 (30 and 100 μM) for 1 h, and then treated with RSL-3 (0.1 μM) for 24 h. Cell viability was assessed by the MTT assay. (C) *DHCR7* expression in various cancer cells was analyzed using the CellExpress system. (D) *DHCR7* mRNA levels in ferroptosis-sensitive cancer cells were assessed. \*\**p* < 0.01 vs. Huh-7 cells. (E) Human hepatocellular carcinoma-derived cell lines were pretreated with Fer-1 (0.5 μM) or AY9944 (30 and 100 μM) for 1 h, and then treated with RSL-3 (0.1 μM) for 24 h. Cell viability was assessed by the MTT assay. (F) *DHCR7* expression in normal tissues and organs was analyzed using the RefEx database. (G) Murine primary hepatocytes (HC) were pretreated with Fer-1 (0.5 μM) or AY9944 (30 and 100 μM) for 1 h, and then treated with RSL-3 (1 μM) for 24 h. Cell viability was assessed by MTT assay. (H) Liver samples and serum were obtained from hepatic ischemia-reperfusion or sham-operated mice treated with AY9944 (25 mg/kg) 1 h prior to open laparotomy. Serum AST and ALT levels, HE staining, and the histopathology score were assessed. Statistical significance was calculated using the one-way ANOVA (D and H) or two-way ANOVA (A, G, and H) with Tukey’s post hoc test. Data are expressed as dot plots and/or means ± SEM. \**p* < 0.05, \*\**p* < 0.01.

### AY9944 prevents ferroptosis of normal hepatocytes in vitro and in vivo

Consistent with its fundamental function as the key enzyme in the synthesis of cholesterol, *DHCR7* expression in the liver is particularly high among normal tissues or organs (RefEx, https://refex.dbcls.jp/about.php?lang=jahttps://refex.dbcls.jp/about.php?lang=ja) (19) (Fig. 4F). Therefore, we investigated the role of DHCR7 in normal hepatocytes. Similar to the results in Huh-7 cells, AY9944 suppressed RSL-3-induced ferroptosis in murine primary hepatocytes (Fig. 4G). We have recently shown that ferroptosis contributes to the development of hepatic IRI (4). Intraperitoneal injection of AY9944 significantly attenuated liver injury, as evaluated by pathological severity scores and serum levels of AST and ALT, after hepatic IRI in mice (Fig. 4H). These findings suggest that DHCR7 is a potential therapeutic target for treating or preventing liver diseases related to ferroptosis.

## Discussion

Ferroptosis is involved in the pathogenesis of many liver diseases and is regarded as a potential target for these disorders; however, the regulatory mechanism of liver ferroptosis is poorly understood (5, 6). In the present study, using the whole-genome screening approach, we identified DHCR7 as a novel regulator of ferroptosis in hepatocytes. DHCR7 inhibition or extrinsic supplementation of DHCR7’s substrate 7-DHC suppressed ferroptosis in Huh-7 cells and normal primary hepatocytes. Furthermore, DHCR7 inhibition ameliorated hepatic IRI *in vivo*. These findings demonstrate that targeting DHCR7 has therapeutic potential for treating and preventing ferroptosis-related diseases.

We clearly demonstrated that 7-DHC is responsible for the suppressive effect of DHCR7 inhibition on hepatic ferroptosis because 7-DHC supplementation, but not cholesterol deprivation, suppressed ferroptosis. 7-DHC is more than just a precursor of cholesterol. For instance, 7-DHC is metabolized to vitamin D by ultraviolet light B in the skin and plays an important role in calcium homeostasis and immune function (13, 14) (Fig. 2B). Previous reports also showed that 7-DHC could be extremely prone to free radical oxidation. In particular, Porter *et al*. (16) reported that the calculated *kp* value of 7-DHC to react with peroxy radicals is about 11 times higher than that of arachidonic acid, oxidation of which is absolutely key for initiating ferroptosis. Indeed, we observed that DHCR7 inhibition increased intracellular levels of 7-DHC and its major oxysterol metabolite, DHCEO, which was further increased by RSL-3 treatment. Therefore, we assume that 7-DHC is rapidly oxidized prior to membrane phospholipid peroxidization, thereby preventing ferroptosis. However, more than 10 kinds of 7-DHC-derived oxysterol metabolites have been identified so far (15): thus, further studies to assess the amounts of each metabolite will clarify the regulatory mechanism of ferroptosis by oxysterol.

Considering the clinical application of ferroptosis regulation, the inhibition of ferroptosis is expected to be a therapeutic option for ferroptosis-related liver diseases, such as hepatic IRI, hemochromatosis, NASH, and acetaminophen-induced acute live failure (5, 6). On the other hand, the induction of ferroptosis is also expected to be a novel therapeutic strategy for intractable diseases including liver fibrosis and cancer because activated hepatic stellate cells and conventional chemotherapy-resistant cancer cells are known to be sensitive to ferroptosis (20, 21). In the present study, we found two different types of ferroptosis-inhibiting strategies: extrinsic 7-DHC supplementation and DHCR7 inhibition. 7-DHC suppressed ferroptosis in a cell-type-independent manner, consistent with its proposed function as a radical scavenger. In contrast, DHCR7 inhibition suppressed ferroptosis, which is probably dependent on high DHCR7 expression. Considering the tissue-specific expression patterns of DHCR7, its inhibition can be a more specific approach for suppressing ferroptosis in hepatocytes. Moreover, it seems likely that DHCR7 inhibition can reduce the side effects on the liver in ferroptosis-inducing therapy against progressive liver fibrosis or chemotherapy-resistant cancers.

An additional point to be noted is that mutations of the *DHCR7* gene in humans cause Smith-Lemli-Opitz syndrome (SLOS), which is characterized by multiple congenital anomalies including microcephaly, developmental delay, typical facial appearance, and cardiac abnormalities (22, 23). The patients show low serum cholesterol levels, and increased serum and tissue 7-DHC levels (24). Although some patients are treated with cholesterol supplementation, bile acid supplementation, or steroid hormone replacement therapy, no effective therapy for SLOS is currently available. Taking the etiology of SLOS into consideration, we cannot rule out the possibility that long-term DHCR7 inhibition can have adverse effects. Furthermore, 7-DHC-derived oxysterol metabolites including DHCEO reportedly have oxidative toxicity that causes retinal degradation in a rat model of SLOS (17). Thus, further investigations are necessary for the development of safe clinical applications to protect against hepatic ferroptosis by DHCR7 inhibition or 7-DHC supplementation.

In conclusion, our results suggest that the terminal enzyme of cholesterol biosynthesis DHCR7 and its substrate 7-DHC suppress hepatic ferroptosis and provide new insights into the regulatory mechanism of liver ferroptosis.

## Acknowledgments

We thank Drs. Tadashi Kasahara (Jichi Medical University), Kazumoto Murata (Jichi Medical University), Hitoshi Endo (Jichi Medical University), Yasushi Saga (Jichi Medical University), Kenji Tago (Jichi Medical University), and Hidetoshi Hayashi (Nagoya City University) for their invaluable suggestions.

## List of Abbreviations

ALD: alcoholic liver disease
ALT: alanine aminotransferase
AST: aspartic aminotransferase
7-DHC: 7-dehydrocholesterol
DHCEO: 3β,5α-dihydroxycholest-7-en-6-one
DHCR7: 7-dehydrocholesterol reductase
Fer-1: Ferrostatin-1
GFP: green fluorescent protein
GPX4: glutathione peroxidase 4
GSH: glutathione
HE: hematoxylin and eosin
IRI: ischemiareperfusion injury
LA: linoleic acid
LC-MS/MS: liquid chromatograph-mass spectrometry
Mβ-CD: Methyl-β-cyclodextrin
NASH: non-alcoholic steatohepatitis
NGS: next-generation sequencer
PUFAs: polyunsaturated fatty acids

