## Supplementary file for "DHCR7 as a novel regulator of ferroptosis in hepatocytes"

*Western blot analysis*

Protein samples were separated by sodium dodecyl sulfate-polyacrylamide electrophoresis (SDS-PAGE) and transferred to PVDF membranes. After blocking with Blocking One (NACALAI TESQUE, Kyoto, Japan) for 30 min, the membranes were incubated overnight at 4 °C with the following primary antibodies for DHCR7 (ab103296; Abcam, Cambridge, UK), GPX4 (sc-177570; Santa Cruz Biotechnology, Dallas, TX, USA), and β-actin (A5441, Sigma; St Louis, MO, USA). HRP-Goat anti mouse Superclonal IgG (Thermo Fisher Scientific; Waltham, MA, USA) or HRP-goat antirabbit IgG (Cell Signaling Technology, Danvers, MA) were used as secondary antibodies and incubated with membrane for 1 h. After washing with TBS-Tween, immunoreactive bands were visualized by Western Blot Quant HRP substrate (TAKARA Bio, Shiga, Japan) or Western BLoT Ultra Sensitive HRP substrate (TAKARA Bio).

*Heteroduplex mobility assay*

The mutations in *DHCR7*-KO cells were assessed by heteroduplex mobility assay (HMA). The genomic DNA was extracted from the cell pellet with phenol-chloroform. The mutated regions of DHCR7 were amplified with the following primers: DHCR7#1 forward, 5′-TTTTAAGCAAATTCACAGGCAAT-3′; DHCR7#1 reverse, 5′-GCCCATTGAAGAACAGCTTG-3′; DHCR7#2 forward, 5′-TTTTAAGCAAATTCACAGGCAAT-3′; DHCR7#2 reverse, 5′-GGAAGGTGACCCACAAGGTA-3′. The amplicon was denatured at 95ºC for 5 min and then gradually cooled in a stepwise manner. The heteroduplex was separated by polyacrylamide gel electrophoresis.

*LC-MS/MS analysis for 7-DHC and DHCEO*

7-DHC analysis: Aliquots of frozen cells were homogenized in PBS. After the lipids were extracted by methanol and chloroform, the chloroform layer was collected and evaporated at 80°C under nitrogen gas. The residue was dissolved in 100 μL of EtOH. 7-dehydroxycholesterol contents was quantified by LC-MS/MS. Briefly, $[^{2}H_{6}]$7-dehydroxycholesterol (5 ng) was added to 2μL of lipids extract as an internal standard, and saponification was carried out in 0.5 mL of 1N ethanolic KOH with butylated hydroxytoluene at 37 °C for 1 h. After the addition of 0.25 mL of distilled water, sterols were extracted with 1 mL of n-hexene, and the extract was evaporated to dryness under a nitrogen gas stream. The reagent mixture for derivatization consisted of 2-methyl-6-nitrobenzoic anhydride (100 mg), 4-dimethylamino- pyridine (30 mg), picolinic acid (80 mg), pyridine (1.5 mL), and triethylamine (200 μL). The freshly prepared reagent mixture (170 μL) was added to the sterol extract, and the reaction mixture was incubated at 80°C for 60 min. After the addition of 1 ml of n-hexane, the mixture was centrifuged at 1,500 g for 5 min. The clear supernatant was collected and evaporated at 80°C under nitrogen gas. The residue was dissolved in 50 μL of acetonitrile, and an aliquot (5 μL) was injected into the following LC-MS/MS system. The LC-MS/MS system consisted of a TSQ Quantum Ultra mass spectrometer (Thermo Fisher Scientific, San Jose, CA) equipped with an H-ESI probe and an Ultimate 3000 HPLC system (Thermo Fisher Scientific, San Jose, CA). Chromatographic separation was performed using a Hypersil GOLD column (150 3 2.1 mm, 3 mm, Thermo Electron) at 40°C, and the following gradient system was used at a flow rate of 300 ml/min: initially, the mobile phase was composed of acetonitrile-methanol-water (40:40:20, v/v/v) containing 0.1% acetic acid; then it was programmed in a linear manner to acetonitrile-methanol-water (45:45:10, v/v/v) containing 0.1% acetic acid over 20 min. The final mobile phase was kept constant for an additional 20 min. The general LC-MS/MS conditions were as follows: spray voltage, 1,000 V; vaporizer temperature, 350°C; sheath gas (nitrogen) pressure, 85 psi; auxiliary gas (nitrogen) flow, 60 arbitrary units; ion transfer capillary temperature, 350°C; collision gas (argon) pressure, 1.5 mTorr; and ion polarity, positive. The multiple ion detector was focused on m/z 553.339 for 7-dehydroxycholesterol, and m/z 560.377 for $[^{2}H_{6}]$7-dehydroxycholesterol was used as an internal standard for 7-dehydroxycholesterol.

DHCEO analysis: The LC-MS/MS system consisted of a 4000 QTRAP tandem mass spectrometer (SCIEX; Tokyo, Japan) equipped with an Exion LC system (SCIEX). Chromatographic separation was performed using an ODS column (5C18-MS-II 5μm, 4.6×250 mm; Nacalai Tesque; Kyoto, Japan) at 40°C. The column was eluted with a mobile phase consisting of solvent A (methanol-water (90:10, v/v) containing 0.1% formic acid) and solvent B (2-propanol containing 0.1% formic acid). The mobile phase gradient profile was as follows: 0–10 min, 0% B; 10–20 min, 0–30% B linear; 20.1 min, 100% B. The flow rate was 1.0 mL/min. The general LC-MS/MS conditions were as follows: declustering potential, 71.0 V; entrance potential, 10.0 V; collision energy, 21.0 V; collision cell exit potential, 18.0 V; temperature, 500°C; source, APCI; and ion polarity, positive. DHCEO was detected by multiple reaction monitoring (MRM) for the transition of precursor ions to products: (*m/z* 399>381). Standard DHCEO (EVU139) was purchased from Kerafast (Boston, MA)

*Supplementary Table 1. sgRNA used for CRISPR/Cas9-mediated genome editing*

| | Forward | Reverse |
|---|---|---|
| sg*DHCR7*#1 | CACCGTCAAACCACTTCCCGATCCG | AAACCGGATCGGGAAGTGGTTTGAC |
| sg*DHCR7*#2 | CACCGTGGGCGGCTTTCCTCGTTAT | AAACATAACGAGGAAAGCCGCCCAC |
| sg*GPX4*#1 | CACCGGTCCCGGGACGCGCACTGCA | AAACTGCAGTGCGCGTCCCGGGACC |
| sg*GPX4*#2 | CACCGTTAACCTGGACAAGTACCGG | AAACCCGGTACTTGTCCAGGTTAAC |

*Supplementary Table 2. Primers used for real-time RT-PCR analysis*

| | Forward | Reverse |
|---|---|---|
| *ACSL4* | CCCTGAAGGATTTGATCTCTTCG | CCTTAGGTCGGCCAGTAGAAC |
| *AIFM2* | GGCCAACATCGTCAACTCTG | ACACCGTCATTTCTCCCCAT |
| *DHCR7* | ATGCCGTCTCCACCTTCG | AACCACTTCCCGATCCGA |
| *DHODH* | CCACGGGAGATGAGCGTTTC | CAGGGAGGTGAAGCGAACA |
| *GPX4* | GCCTTCCCGTGTAACCAGT | GCGAACTCTTTGATCTCTTCG |
| *SLC7A11* | ATGCAGTGGCAGTGACCTTT | GGCAACAAAGATCGGACTG |
| *ACTB* | GGCACTCTTCCAGCCTTCCTTC | GCGGATGTCCACGTCACACTTCA |

**Supplementary figure legends**

*Supplementary Fig. S1. DHCR7-KO cells*

*DHCR7*-KO Huh-7 cells were generated using the CRISPR/Cas9 genome editing system. (A) The indel mutations of each sgRNA analyzed by the HMA method. (B) The cells were treated with pitavastatin (PTV, 1 mM) for 24 h to enhance DHCR7 expression and the effectiveness of *DHCR7*-KO was analyzed by western blotting.

*Supplementary Fig. S2. Quantitative analysis of C11-BODIPY$^{581/591}$ fluorescence intensity*

Control (sgGFP) and *DHCR7*-KO (sgDHCR7) Huh-7 cells were treated with or without RSL-3 (0.1 μM) for 6 h. The fluorescence intensity of C11-BODIPY$^{581/591}$ was measured at 30 min intervals (n =3).

A

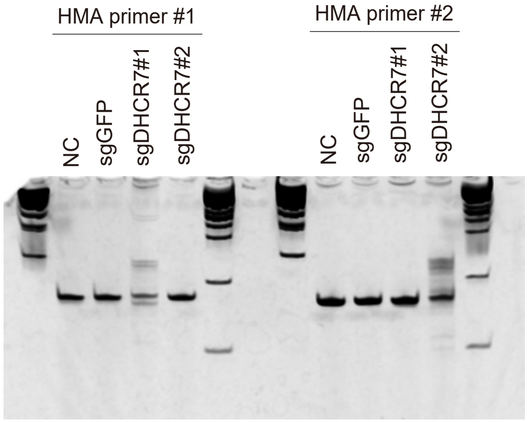
HMA primer #1
HMA primer #2
NC
sgGFP
sgDHCR7#1
sgDHCR7#2
NC
sgGFP
sgDHCR7#1
sgDHCR7#2

B

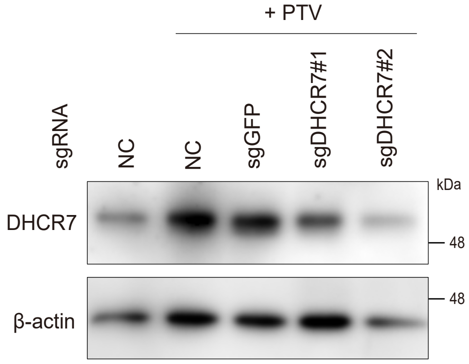
+ PTV
sgRNA
NC
NC
sgGFP
sgDHCR7#1
sgDHCR7#2
kDa
DHCR7
48
β-actin
48

A

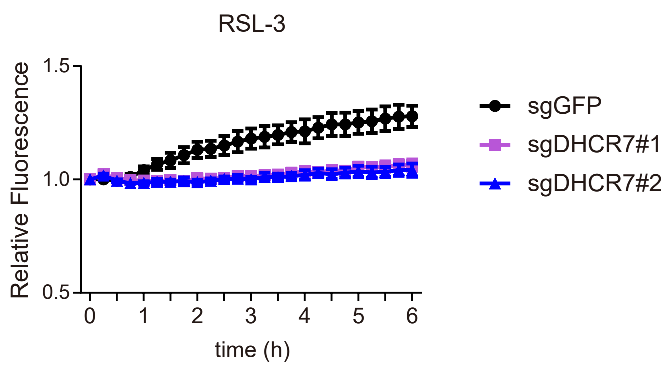
RSL-3
Relative Fluorescence
1.5
1.0
0.5
0
1
2
3
4
5
6
time (h)
sgGFP
sgDHCR7#1
sgDHCR7#2

B

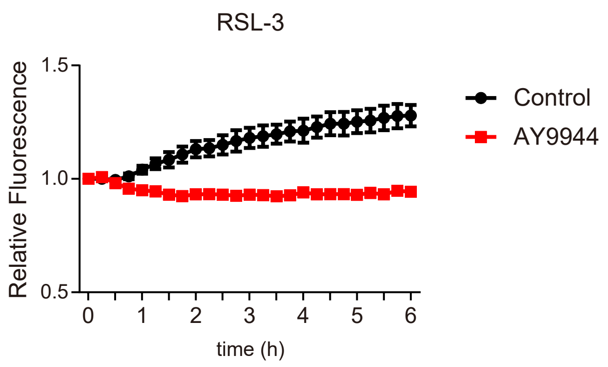
RSL-3
Relative Fluorescence
1.5
1.0
0.5
0
1
2
3
4
5
6
time (h)
Control
AY9944
